# Enhancement of the Activity of the Antimicrobial Peptides HNP1 and LL-37 by Bovine Pancreatic Ribonuclease A

**DOI:** 10.1101/2022.06.16.496515

**Authors:** Bryan Ericksen

## Abstract

**Background:** HNP1, LL-37, and HBD1 are antimicrobial against *Escherichia coli* ATCC 25922 at the standard inoculum but less active at higher inocula.

**Methods:** The virtual colony count (VCC) microbiological assay was adapted for high inocula and the addition of yeast tRNA and bovine pancreatic ribonuclease A (RNase). 96-well plates were read for 12 hours in a Tecan Infinite M1000 plate reader and photographed under 10x magnification.

**Results:** Adding tRNA 1:1 to HNP1 at the standard inoculum almost completely abrogated activity. Adding RNase 1:1 to HNP1 at the standard inoculum of 5x10^5^ CFU/mL did not enhance activity. Increasing the inoculum to 6.25x10^7^ CFU/mL almost abrogated HNP1 activity. However, adding RNase 25:1 to HNP1 enhanced activity. Adding both tRNA and RNase resulted in enhanced activity, indicating that the enhancement effect of RNase overwhelms the inhibiting effect of tRNA when both are present. HBD1 activity at the standard inoculum was almost completely abrogated by the addition of tRNA, but LL-37 activity was only slightly inhibited by tRNA. At the high inoculum, LL-37 activity was enhanced by RNase. HBD1 activity was not enhanced by RNase. RNase was not antimicrobial in the absence of antimicrobial peptides. Cell clumps were observed at the high inoculum in the presence of all three antimicrobial peptides and at the standard inoculum in the presence of HNP1+tRNA.

**Conclusions:** Antimicrobial peptide-ribonuclease combinations have the potential to be active against high cell concentrations and biofilms, conditions where the antimicrobial agent alone is relatively ineffective.

## Introduction

Although cationic antimicrobial peptides (CAPs) have been studied as possible therapeutic agents for many years, few have survived clinical trials to become useful antibiotics. (Mishra 2017) Three CAPs are representative of three different structural classes that contribute to the human innate immune system: human neutrophil peptide 1 (HNP1), an alpha defensin; human beta defensin 1 (HBD1); and the human cathelicidin LL-37. (De Smet 2005) One reason why CAP drug candidates have failed to gain approval is a lack of efficacy. (Magana 2020) I demonstrated a pronounced inoculum effect when the defensin HNP1 was assayed against high inocula of *Escherichia coli* ATCC 25922, such that the antimicrobial peptide almost completely lost activity under those conditions. (Ericksen 2020) A pronounced inoculum effect was also observed when HNP1 was assayed against *Staphylococcus aureus* ATCC 29213 and *Bacillus cereus* ATCC 10876. What might cause this decrease in efficacy at high cell concentrations? The molecular basis of the inoculum effect is unclear. However, one possibility is that bacterial cells produce defensin inhibitors that are at a higher concentration when there are more cells present.

One possible type of inhibition is that polyanions might bind and inhibit CAPs by electrostatic attraction. Here I hypothesize that the polyanion tRNA might inhibit CAPs, that inhibition by RNA is partially responsible for the inoculum effect, and that the addition of ribonuclease could enhance antimicrobial peptide activity, restoring some of the efficacy lost at high cell concentrations.

## Methods

The VCC assay was adapted for high inocula as previously described (Ericksen 2020), and volumes were adjusted to allow for the addition of yeast tRNA (Sigma from *Saccharomyces cerevisiae*) and/or bovine pancreatic ribonuclease (Roche or Macherey-Nagel (MN)). HNP1, LL-37 and HBD1 were synthesized ABI 433A synthesizer using an optimized HBTU activation/DIEA in situ neutralization protocol developed by Kent and coworkers for Boc chemistry solid phase peptide synthesis as previously described. (Zhao 2013)(Pazgier 2013)(Bharucha 2021) Two inocula were studied: the standard inoculum of 5x10^5^ CFU/mL, with cells from a seed culture diluted in 10 mM sodium phosphate pH 7.4, and a high inoculum of 6.25x10^7^ CFU/mL, equivalent to adding undiluted seed culture. Antimicrobial peptides were incubated in 10 mM sodium phosphate pH 7.4 plus 1% tryptic soy broth (TSB) for two hours at 37°C shaking every 5 minutes for 3 seconds in a Tecan Infinite M1000 plate reader. An equal volume of twice-concentrated Mueller Hinton Broth was then added and 96-well plates were read for 12 hours in the plate reader and then some wells containing cell clumps were photographed under 10x magnification. In one experiment, the concentration of TSB present in phosphate buffer was adjusted.

## Results

Adding tRNA 1:1 to HNP1 at the standard inoculum almost completely abrogated activity (Figure 1). Adding Roche RNase 1:1 to HNP1 at the standard inoculum of 5x10^5^ CFU/mL did not enhance activity. Increasing the inoculum to 6.25x10^7^ CFU/mL almost abrogated HNP1 activity. (Figure 2) However, adding RNase 25:1 to HNP1 enhanced activity at the high inoculum. Adding both tRNA and RNase resulted in enhanced activity, indicating that the enhancement effect of RNase overwhelms the inhibiting effect of tRNA when both are present. HBD1 activity at the standard inoculum was almost completely abrogated by the addition of tRNA, but LL-37 activity was only slightly inhibited by tRNA. (Figure 3) At the high inoculum, LL-37 was enhanced, but LL-37 showed greater activity than HNP1 in the absence of RNase. (Figure 4) HBD1 activity was not enhanced by RNase. RNase was not active in the absence of antimicrobial peptides. The observations with HNP1 at the high inolculum were repeated using a second RNase manufacturer, Macherey-Nagel. (Figure 5) The experiment with MN RNase was repeated. (Figure 6) 1% TSB was used in most assays, but the %TSB was varied in one experiment, resulting in maximum activity at 4% TSB with either 5x or 25x MN RNase added. (Figure 7) Cell clumps were observed at the high inoculum in the presence of all three antimicrobial peptides with or without RNase and at the standard inoculum in the presence of HNP1+tRNA. (Figure 8) The VCC assays were conducted with TSB added to the 10 mM sodium phosphate incubation buffer. The enhancement of activity caused by RNase was observed with LL-37 but not HNP1 when washed cells were used, indicating that RNase operates by different mechanisms with the two antimicrobial peptides. Although biofilm formation was not directly assayed, it is assumed that the cell clumps photographed at 10x magnification are biofilms. Ribonuclease did not enhance HBD1 activity at the 6.25x10^7^ CFU/mL inoculum, demonstrating a strong inoculum effect with HBD1 vs. *E. coli*. LL-37 had a much lesser inoculum effect against *E. coli*. The effect of ribonuclease on HNP1 is strongest with lowest amounts of TSB present in the phosphate buffer during the 2 hour incubation. The ability of tRNA to abrogate HNP1 and HBD1 activity, and the failure of tRNA to affect LL-37 activity, at the standard inoculum cannot be explained by net charge. Possibly, hydrophobic interactions play a role in tRNA binding and inhibition. It is also possible that tRNA inducing biofilm formation impacts HNP1 and HBD1 more than LL-37.

**Figure 1.**
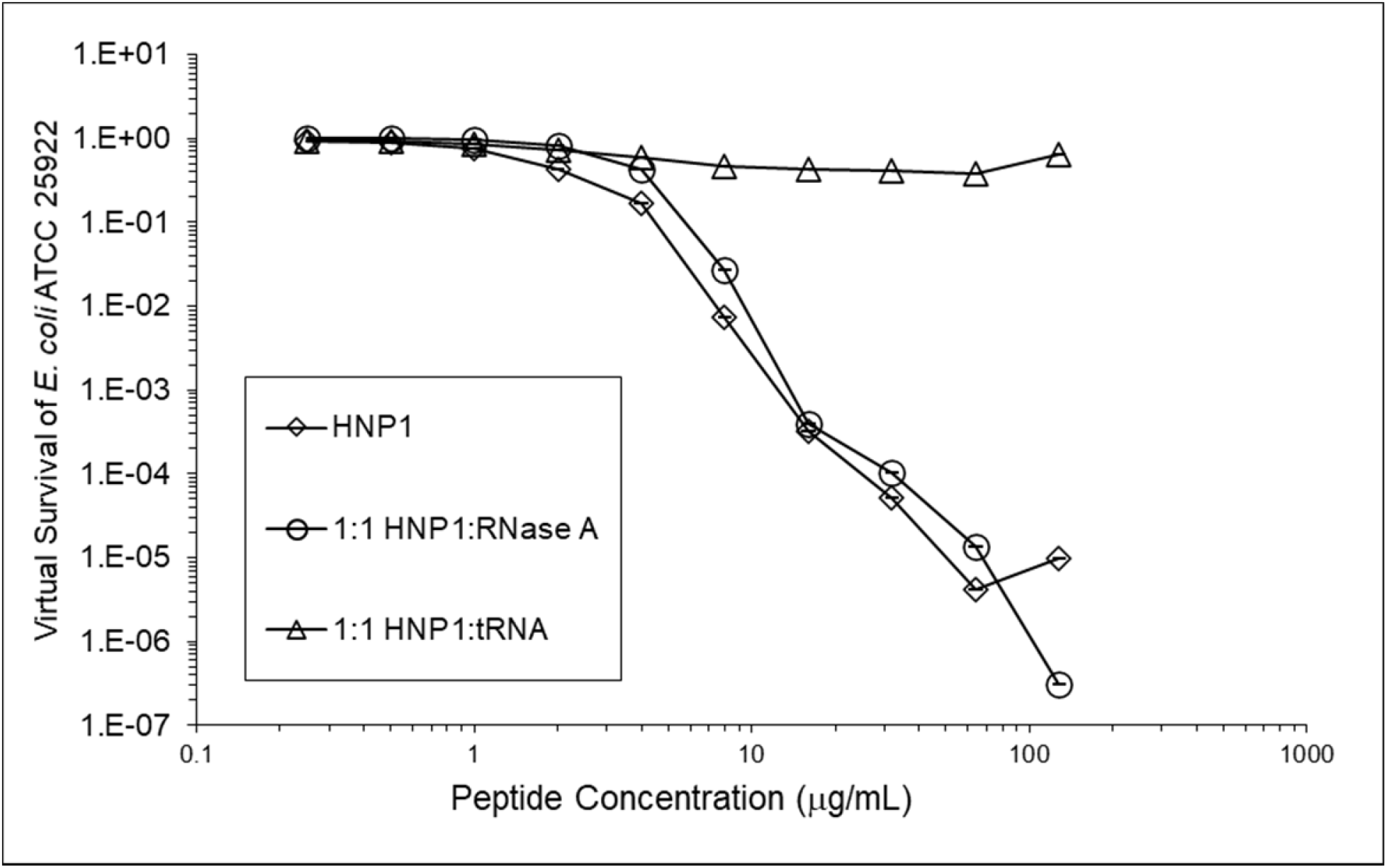
Activity of HNP1 with or without tRNA and RNase at the standard inoculum.

**Figure 2.**
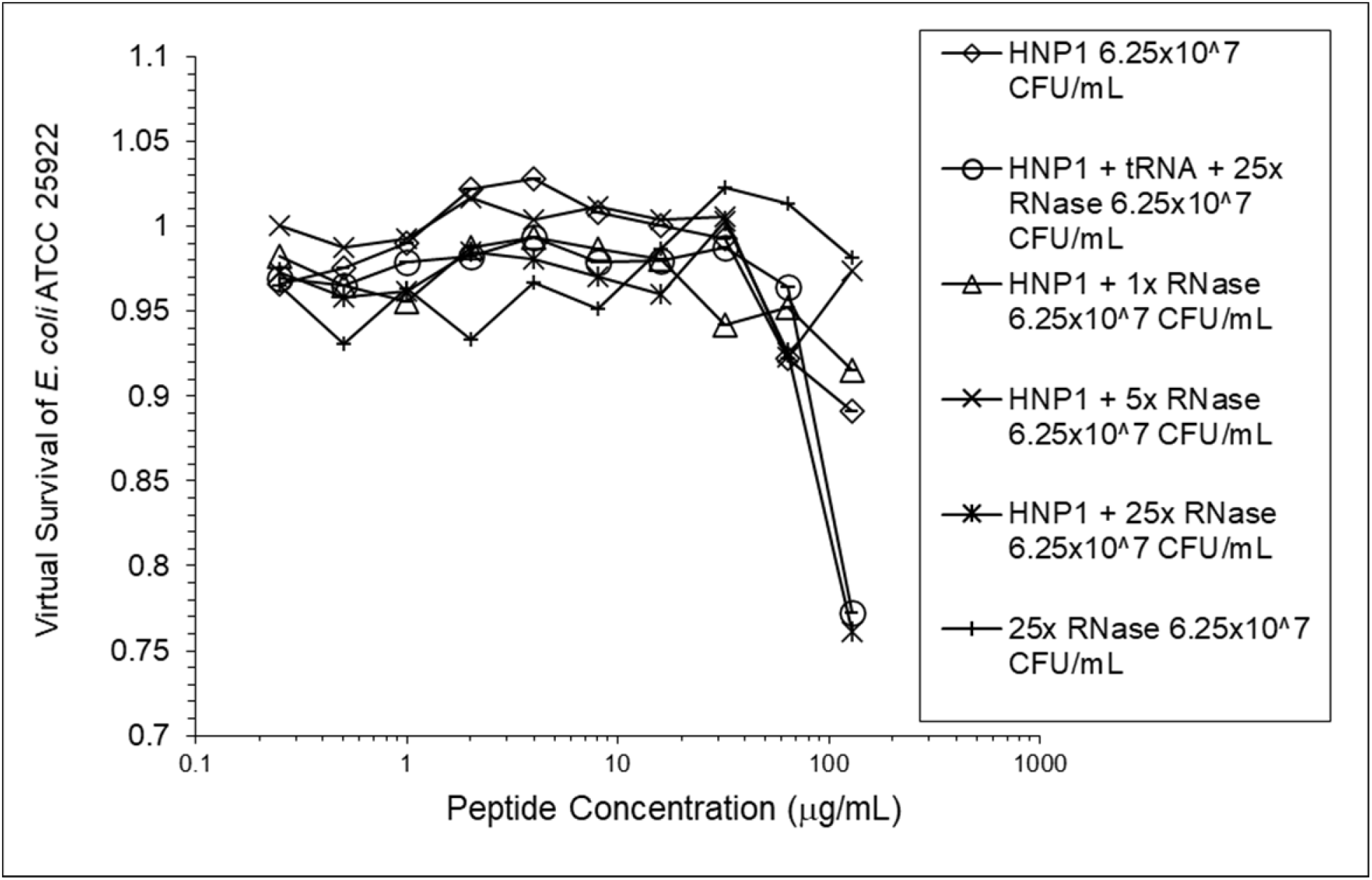
Activity of HNP1 at the high inoculum with or without tRNA and three concentrations of RNase. Activity with HNP1 and both tRNA and the highest concentration of RNase was essentially the same as HNP1 plus RNase alone, indicating the enhancement of activity overcomes inhibition by tRNA. RNase in the absence of antimicrobial peptides was not antimicrobial.

**Figure 3.**
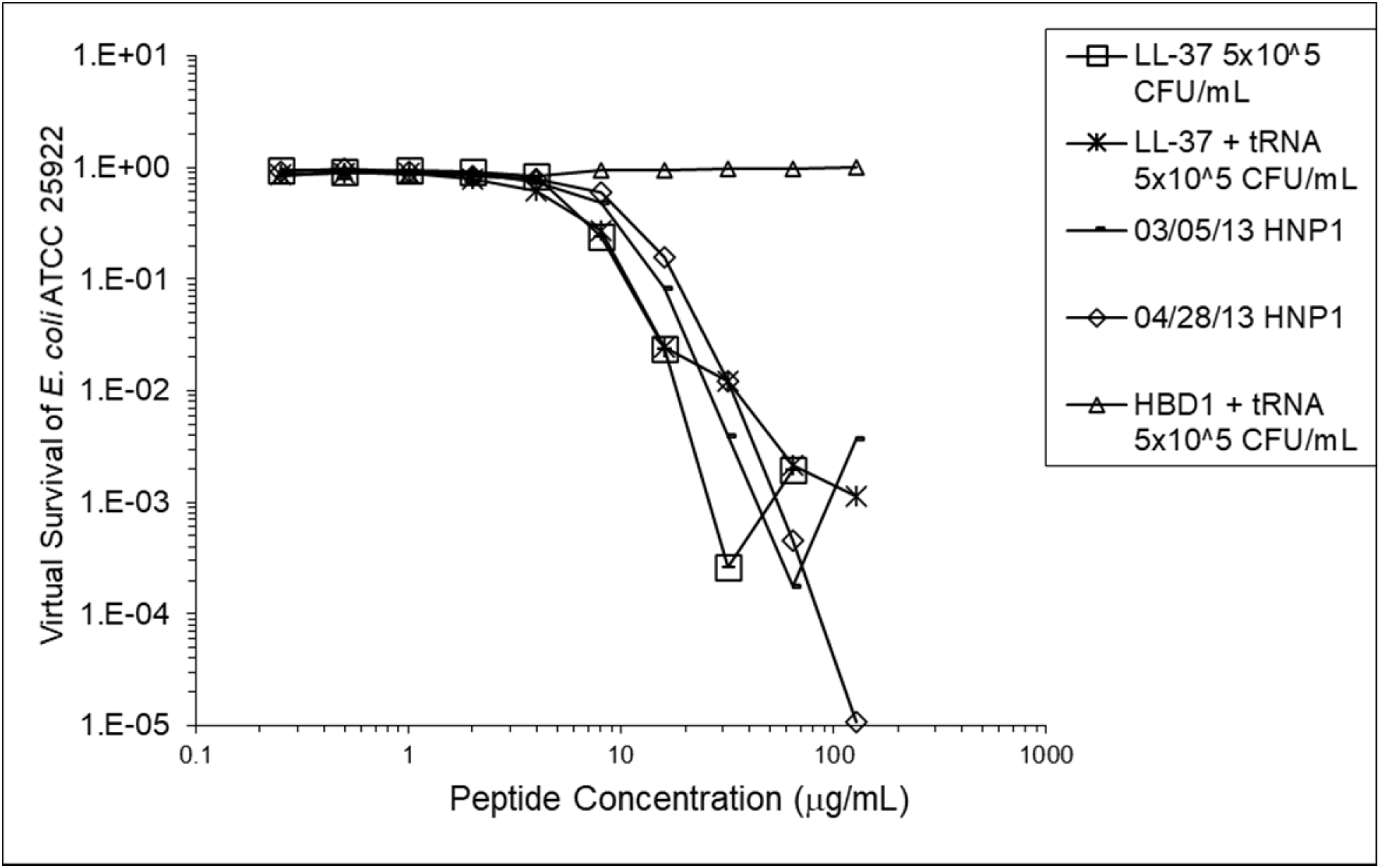
LL-37 was assayed at the standard inoculum with or without tRNA. HBD1 was assayed at the standard inoculum in the presence of 1:1 tRNA. Two preparations of HNP1 were assayed in the absence of tRNA as positive controls.

**Figure 4.**
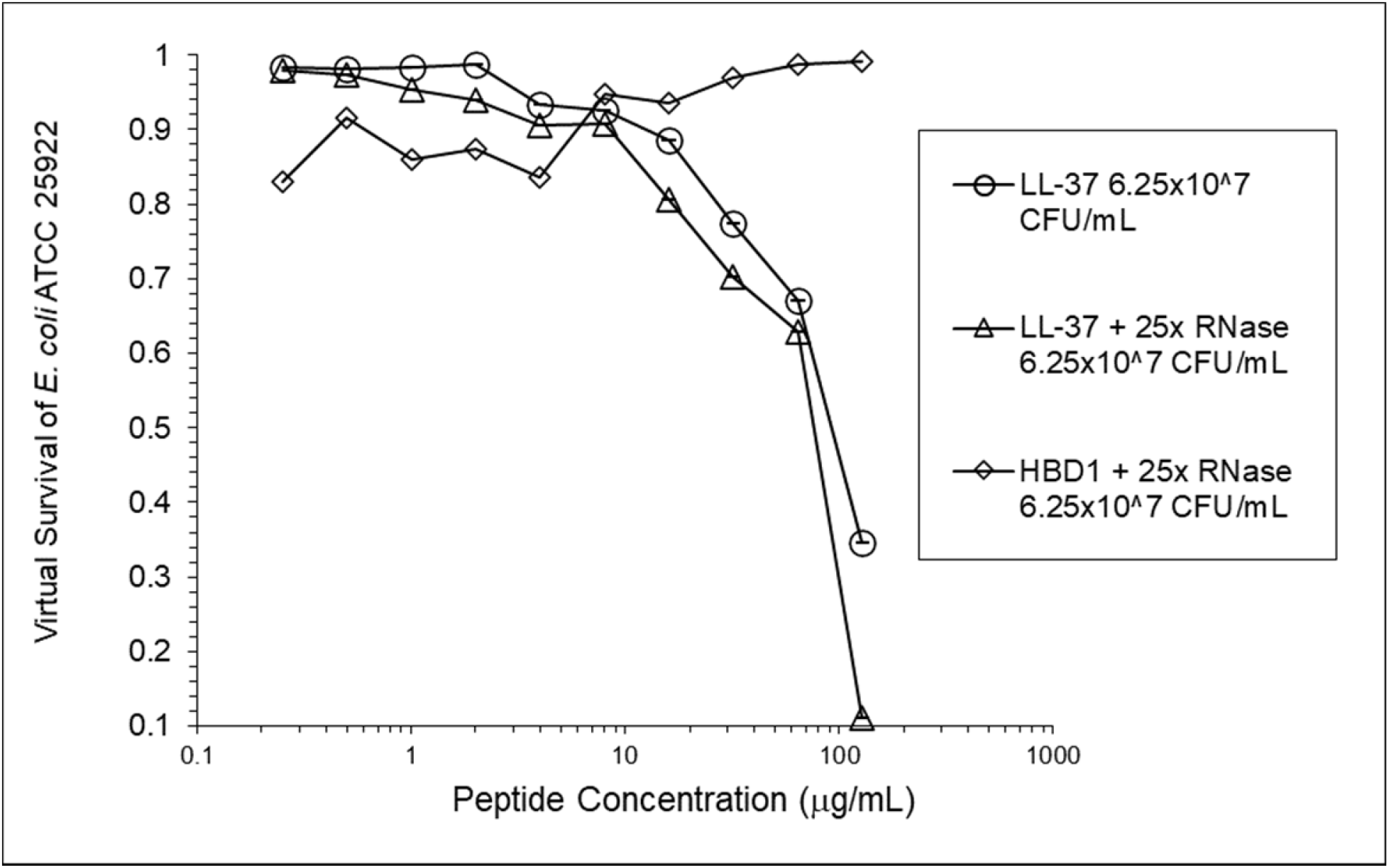
LL-37 was assayed at the high inoculum with or without RNase. HBD1 was assayed at the high inoculum with RNase.

**Figure 5.**
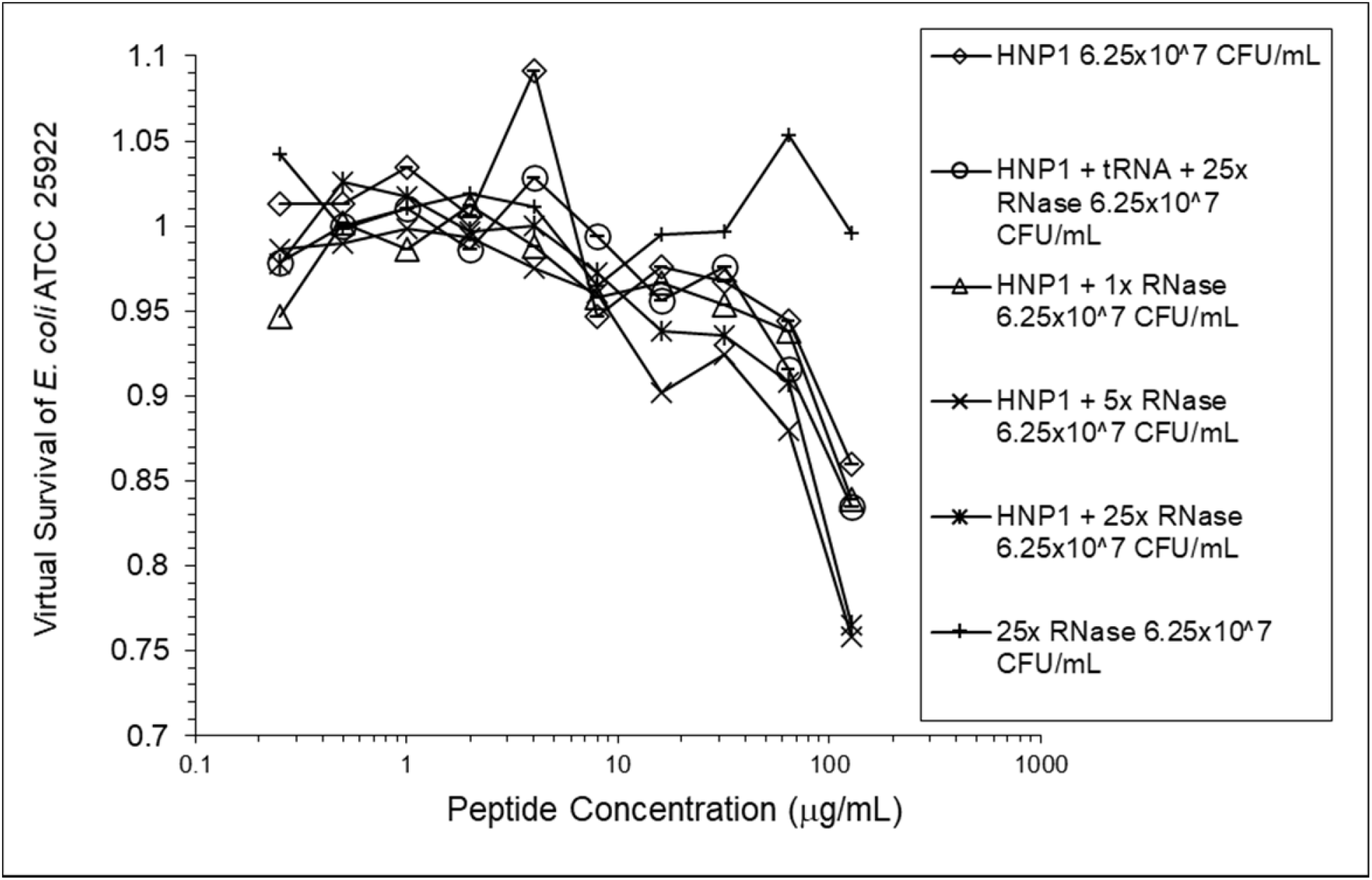
HNP1 was assayed at the high inoculum in the presence and absence of RNase from a second manufacturer, and in the presence of both tRNA and RNase.

**Figure 6.**
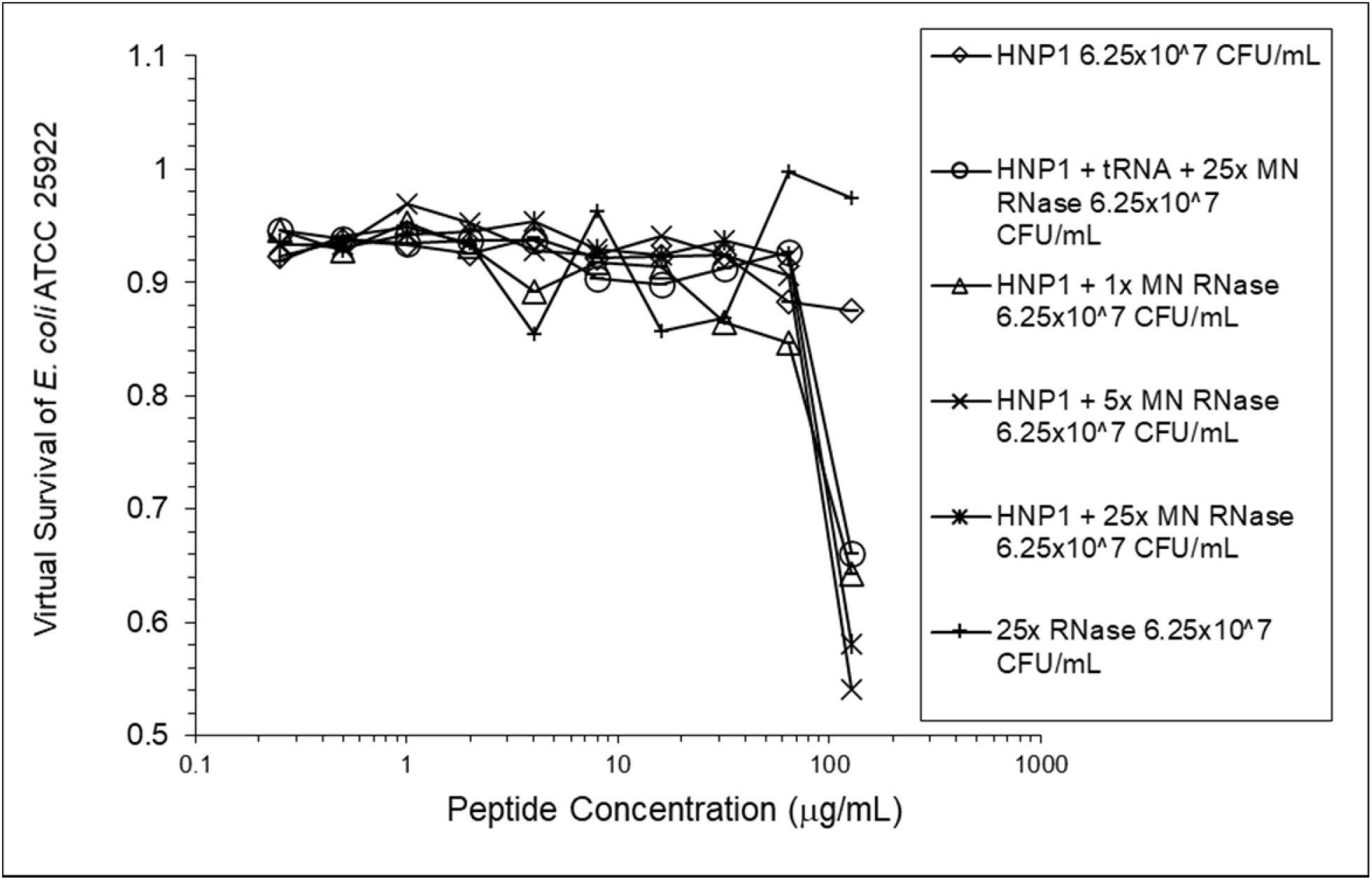
The assay shown in Figure 5 was repeated.

**Figure 7.**
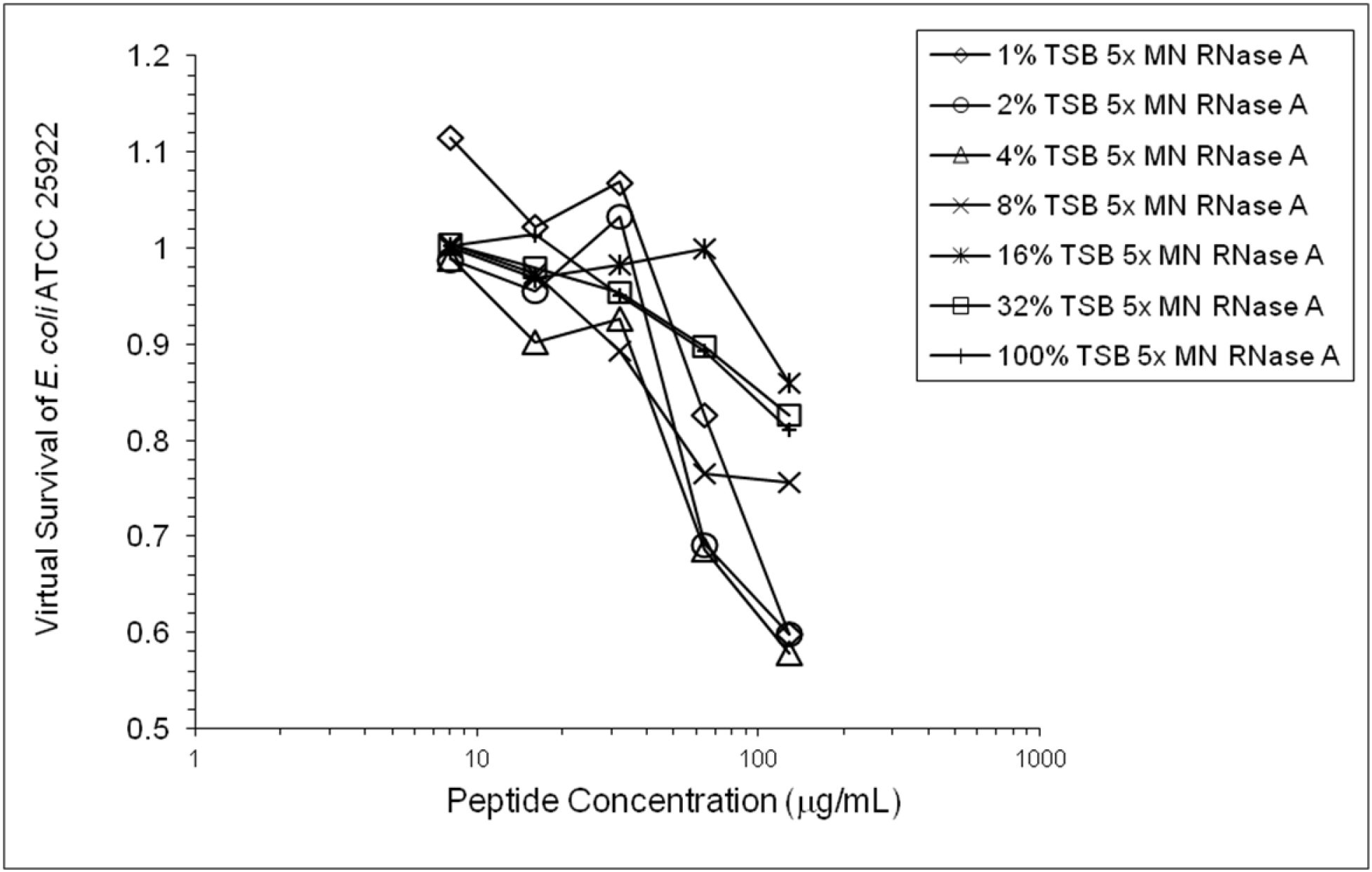
HNP1 was assayed at the high inoculum with variation in the amount of TSB present during the two-hour incubation in 10 mM sodium phosphate buffer.

**Figure 8.**
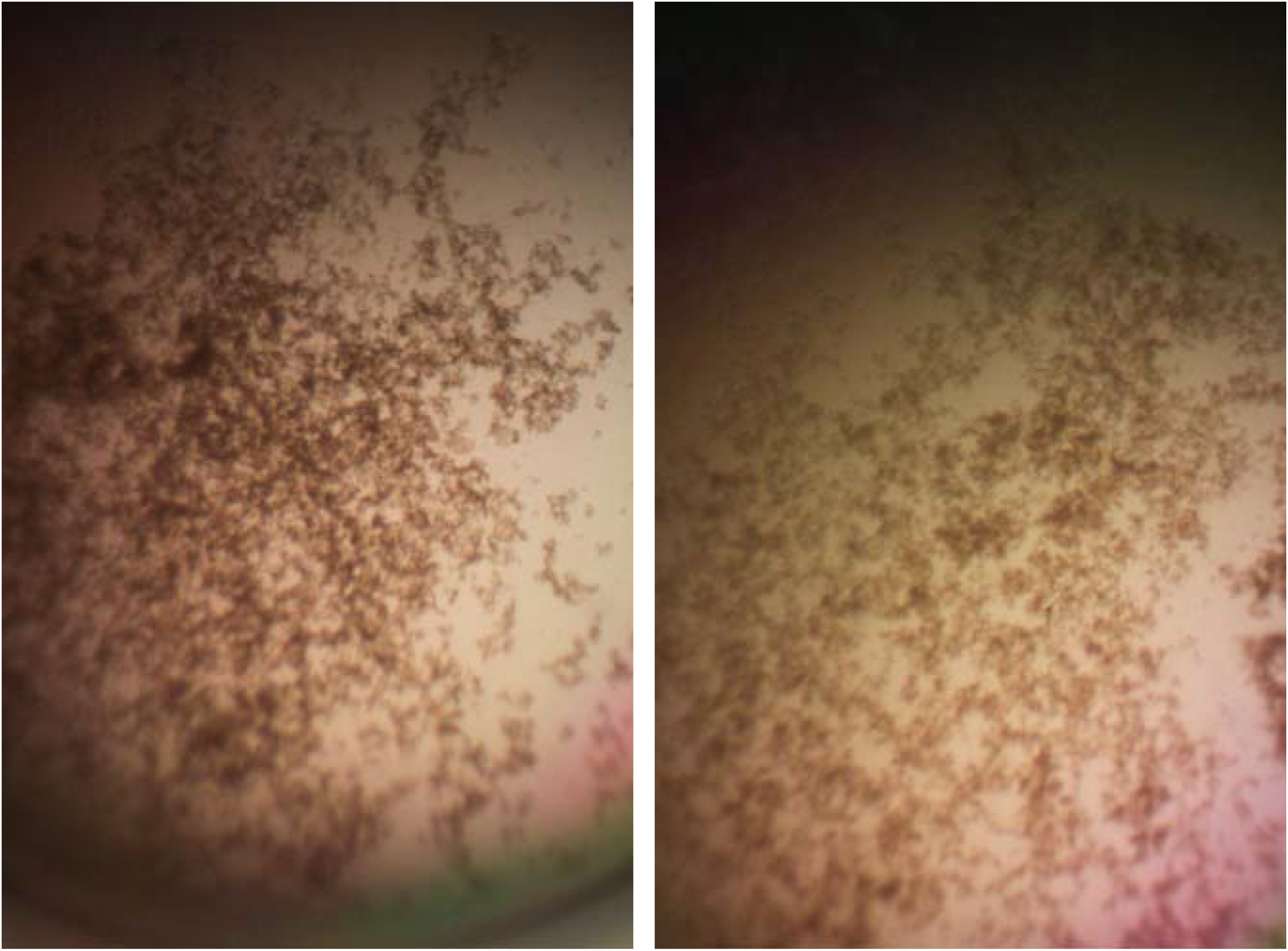
Cell clumps photographed at 10x magnification. Left panel: 128 µg/mL HNP1 at the high inoculum. Right panel: 128 µg/mL HNP1 + 1:5 RNase at the high inoculum.

## Conclusions

Antimicrobial assays are ordinarily conducted using a single antimicrobial agent, studying its effect in isolation. However, the experiments presented here may offer a glimpse into a more realistic in vivo scenario, in which multiple antimicrobial agents work in concert against infection. Eight RNases are encoded by the human genome, many of which have potent antimicrobial activity, such as RNase 7 expressed in epithelial cells. (Sorrentino 2010) Bovine pancreatic RNase A, on the other hand, has a digestive function degrading RNA and an antimicrobial function has not normally been ascribed to it. RNase A is a basic protein (pI = 9.63). It is unknown whether the RNA-degrading activity of RNase or its cationicity is responsible for the enhancement of HNP1 and LL-37 activity. Product literature suggests assaying RNase A using 100 mM Tris buffer, pH 7.4. Enzymatic activity in 10 mM sodium phosphate buffer was not tested, but RNase A is very stable with four disulfide bonds.

The variation in the amount of TSB present in 10 mM phosphate buffer revealed that the increase in activity caused by a small amount of nutrients present, allowing some growth during the two-hour incubation, is counterbalanced by the inhibition of defensin activity at higher TSB concentrations, presumably by the salt content of TSB. This same effect is probably partially responsible for the almost complete abrogation of activity of HNP1 when undiluted seed culture is added to the 96-well plate at the high inoculum in the absence of RNase, since the salt concentration is much higher than in assays at the standard inoculum where the seed culture is diluted in 10 mM sodium phosphate buffer before adding to the 96-well plate.

The vast majority of published VCC assays were conducted at the standard inoculum, reflecting a general reliance on the standard inoculum in a wide range of published antimicrobial assays. Under these conditions, cells are predominantly planktonic. However, a high inoculum may be more medically relevant, since high cell concentrations and biofilms can accompany acute infections. This study demonstrates the utility of conducting assays at a high inoculum, revealing details of antimicrobial activity that would be missed if the antimicrobial agents were studied only at the standard inoculum. Further studies using animal models are necessary to determine whether the enhancement of activity observed at the high inoculum is sufficient to enable the infected host to overcome bacterial infections.

It should be emphasized that both RNA and ribonucleases are ubiquitous in vivo. Therefore, these experiments may be more biologically relevant than VCC experiments lacking RNA or ribonuclease. There are several possible sources of bacterial RNA that might be present at the site of a bacterial infection. Firstly, bacteria normally secrete RNA during their growth, which may have a role in the extracellular matrix of biofilms. (Ozoline 2019) The results of the experiments presented here suggest that this secreted RNA may also be a bacterial defense mechanism against antimicrobial peptides. Secondly, once antimicrobial peptides are released at the infection site, cell lysis may result in the release of intracellular RNAs, including mRNA and tRNA. Thirdly, host RNA may be present. Therefore, inhibition by RNA must be regarded as a common obstacle to effective antimicrobial activity that frequently occurs in real world scenarios.

The combination of an antimicrobial peptide with a ribonuclease could be regarded as a novel invention that could possibly be used as a therapy to treat bacterial infections. LL-37 and RNase 1 have been shown to act synergistically to kill *E. coli*. (Eller 2020) RNases have been tested in clinical trials as chemotherapeutics for the treatment of cancer. (Ardelt 2009)

Further studies are warranted to determine whether these results could be generalized to antimicrobial peptide-nuclease combinations, as might be suggested by the presence of DNA in biofilms. A combination of an antimicrobial peptide with both deoxyribonuclease (DNase) and RNase might be expected to be more potent than the combination of the antimicrobial peptide and RNase in the absence of DNase, because DNA is considered a more prevalent structural component of biofilms than RNA. (Gilan 2013) DNase is an approved drug, dornase alfa (Pulmozyme), which cuts apart extracellular DNA in the lungs of cystic fibrosis patients, making the mucus thinner and easier to expel. (Wagener 2012) It is possible that DNase in combination with an antimicrobial peptide and RNase would form an effective treatment against acute bacterial infections. A new generation of antimicrobial peptide-nuclease combinations would offer a new hope that peptides that are sometimes defeated by the resistance mechanism of biofilm formation can be repurposed to degrade biofilms instead, with increased activity to fight infections.

## Acknowledgments

I thank Robert I. Lehrer for sharing an unpublished manuscript proposing that defensins are inhibited by RNA, Wuyuan Lu for providing antimicrobial peptides and helpful discussions, and Peprotech, Inc. for funding.

